# LIPT1 loss confers replication stress and PARP inhibitor sensitivity through PrimPol-mediated ssDNA gaps

**DOI:** 10.1101/2025.09.30.679512

**Authors:** Zengfu Shang, Jui-Chung Chiang, Ching-Cheng Hsu, Ciara Newman, Anthony J Davis, Yuanyuan Zhang

**Affiliations:** Department of Radiation Oncology, University of Texas Southwestern Medical Center, Dallas, TX, USA; Harold C. Simmons Comprehensive Cancer Center, University of Texas Southwestern Medical Center, Dallas, TX, USA

## Abstract

Replication stress (RS) and altered metabolism are two hallmarks of cancer, yet how metabolic perturbations contribute to RS remains poorly understood. Lipotransferase 1 (LIPT1) catalyzes the covalent attachment of lipoic acid to mitochondrial 2-ketoacid dehydrogenases, sustaining flux through the tricarboxylic acid (TCA) cycle. Loss of LIPT1 causes accumulation of 2-hydroxyglutarate (2-HG), which is known to inhibit α-ketoglutarate (α-KG)–dependent histone demethylases and promotes heterochromatin formation. Here, we show that 2-HG–driven heterochromatin impedes replication fork progression, causing fork stalling and RS in LIPT1-deficient cancer cells. To bypass stalled forks, PrimPol-mediated repriming resumes DNA synthesis but leaves behind single-stranded DNA (ssDNA), which requires poly (ADP-ribose) polymerase 1 (PARP1) for repair. Furthermore, nascent DNA at reprimed forks undergoes MRE11-dependent degradation, further destabilizing replication fork integrity. Consequently, LIPT1 deficiency promotes replication and genome instability, and therapeutic vulnerability to PARP inhibitor. Together, these findings reveal a mechanistic link between mitochondrial lipoylation and replication fork stability, uncovering a metabolic basis for genome instability in cancer.

## Introduction

DNA replication is essential for cell survival and faithful transmission of genetic information. However, replication forks often encounter endogenous and exogenous obstacles that can stall their progression, a phenomenon broadly described as replication stress (RS) (*1*). Elevated RS is a hallmark of cancer, as oncogenes accelerate DNA synthesis, induce RS and genomic instability during early carcinogenesis (*2, 3*). To tolerate chronic RS, cancer cells rely on DNA damage tolerance (DDT) and repair mechanisms, which present vulnerability to therapeutically target (*4, 5*).

DNA replication is also metabolically demanding, requiring energy and metabolites as biosynthetic precursors and cofactors (*6-8*). The influence of nutrient availability on replication has long been recognized in bacteria, yeast, and mammalian cells (*6, 7*), yet the molecular mechanisms linking metabolism to genome replication remains incompletely defined. Emerging evidence suggests that central carbon metabolites and metabolic enzymes directly impact replication functions (*9, 10*). For instance, nuclear ATP-citrate lyase, succinate dehydrogenase (SDH) and pyruvate dehydrogenase (PDH) complexes contribute to histone modifications (*11-13*). The TCA cycle metabolite alpha-ketoglutarate (α-KG) serves as a cofactor for Jumonji-domain histone lysine demethylases and other epigenetic regulators, thereby regulating chromatin methylation (*14-16*). In contrast, oncometabolite 2-hydroxyglutarate (2-HG), due to structural similarity to α-KG, competitively inhibits these enzymes, altering chromatin structure and replication dynamics (*17, 18*). In addition, glycolytic enzymes such as phosphoglycerate kinase (PGK), lactate dehydrogenase (LDH), and glyceraldehyde 3-phosphate dehydrogenase (GAPDH) directly interact with replication machinery and modulate replicative polymerases (*19, 20*).

Lipoylation, the covalent attachment of an essential lipoic acid (LA) to enzymatic complexes like α-KG dehydrogenase and PDH, is catalyzed by Lipoyltransferase 1 (LIPT1). This modification is essential for central carbon metabolism (*21, 22*). Loss-of-function mutations in LIPT1 causes a class of inborn error of metabolism characterized by metabolic perturbation, neurodevelopmental delay, and childhood mortality (*23-25*). Introducing human LIPT1 mutation into mice results in embryonic lethality (*21*). Beyond its metabolic role, lipoylation has gained attention in cancer biology due to its involvement in cuproptosis, a copper-dependent form of regulated cell death (*26-29*). Utilizing CRISPR screen, our group identified that LIPT1 is a critical metabolic determinant of radiation response in lung cancer cells. Mechanistically, loss of lipoylation leads to 2-HG accumulation, which disrupts H3K9 trimethylation at DNA damage sites, impairs TIP60 recruitment, and compromises ATM activation and homologous recombination repair (*30*).

Here, we report that LIPT1 safeguards DNA replication and genome stability. LIPT1 deficiency induces RS via 2-HG–driven, HP1-dependent chromatin compaction, which impedes replication fork progression. To overcome these obstacles, LIPT1-deficient cells rely on PrimPol-mediated repriming, which generates single-stranded DNA (ssDNA) gaps and nascent strands vulnerable to MRE11-dependent degradation. This combination of gap accumulation, strand degradation, and defective DNA damage repair ultimately drives genomic instability. Collectively, our findings establish LIPT1 as a central regulator linking mitochondrial metabolism to chromatin dynamics, replication fidelity, and genome integrity.

## Results

### LIPT1-deficient cells exhibit slower proliferation and replication fork progression

We first compared replication rate of *LIPT1^-/-^* H460, H157 and HeLa cells with their wild-type (WT) parental controls. As previously reported (*21, 30*), LIPT1 loss reduced lipoylation of DLAT and DLST, which was restored by stable expression of exogenous Myc-tagged LIPT1 (Fig. 1A-C). Across all three cell lines, LIPT1 loss led to slower cellular proliferation (Fig. 1D–F) and reduced colony formation (Fig. 1G–J).

**Fig. 1.**
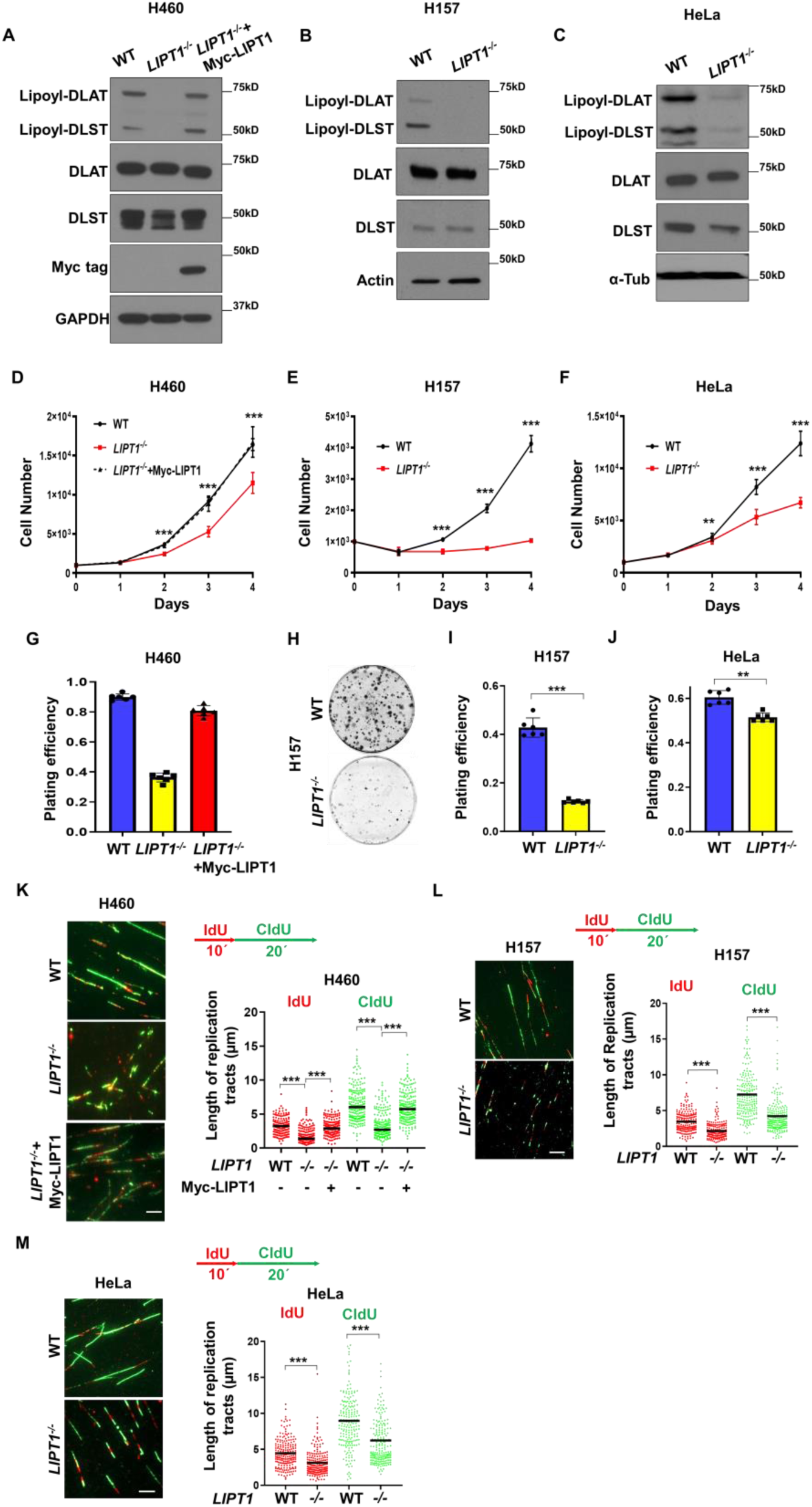
*LIPT1^-/-^*cells exhibit slower proliferation and DNA replication. (**A-C**) Immunoblots of total and lipoylated DLAT and DLST (E2 subunits of PDH and α-KGDH, respectively) in indicated cell lines. GAPDH, Actin and α-Tubulin serve as a loading control. (**D-F**) Cell proliferation of the indicated WT and *LIPT1^-/-^*cell lines. Data are presented as mean ± SD from 8 independent experiments. (**G-J**) Colony formation assays of WT and *LIPT1*^-/-^ H460 cells (G), H157 cells (H-I) and HeLa cells (J). Plating efficiency is shown as mean ± SD from 6 independent experiments. (**K-M**) DNA fiber assay of the indicated cells sequentially pulse-labeled with iododeoxyuridine (IdU, 10 min) and chlorodeoxyuridine (CldU, 20 min). Representative images (left) and quantification of IdU (red) and CldU (green) tract lengths (right) are shown. Scale bar, 10 μm. More than 200 replication tracts were analyzed per condition from 3 independent experiments. The horizontal bars represent the mean of each group. Wilcoxon rank-sum test was used for comparisons between two groups, while two-way ANOVA test was applied for comparisons among more than two independent groups. Statistical significance is indicated as, \*\**P* < 0.01 and \*\*\**P* < 0.001.

We next assessed DNA synthesis by assessing the incorporation of 5-ethynyl-2′-deoxyuridine (EdU). *LIPT1^-/-^* H460 and HeLa cells incorporated less EdU than WT controls, indicating impaired DNA synthesis (Fig. S1A-B). To determine whether this defect was due to reduced replication fork speed, we performed DNA fiber analysis after sequential pulse-labeling cells with iododeoxyuridine (IdU) and chlorodeoxyuridine (CIdU) in WT and *LIPT1^-/-^* cells. Individual fiber measurements revealed a reduction in fork progression speed in *LIPT1^-/-^* cells compared with WT controls (Fig. 1K-M). Consistently, treatment of WT HeLa cells with CPI-613, a lipoic acid analog that inhibits lipoylation, also slowed replication fork progression (Fig. S1C). Together, these findings demonstrate that lipoylation sustain normal replication fork progression and cellular proliferation.

### 2-HG-mediated heterochromatin impedes replication fork in LIPT1-deficient cells

We and others have reported that LIPT1 deficiency causes accumulation of the oncometabolite 2-hydroxyglutarate (2-HG) in cancer cells and in cells and plasma of patients with LIPT1 mutation (*21, 30*). 2-HG promotes histone methylation by inhibiting a-KG dependent demethylases, which have been linked to reduced replication fork progression (*17, 18, 30, 31*). Consistent with this mechanism, we found that the facultative heterochromatin mark H3K27me3 was elevated in *LIPT1^-/-^* H460 and H157 cells, and this increase was fully reversed by α-KG or dimethyl-α-KG treatment (Fig. 2A and C). Notably, α-KG supplementation restored replication fork progression in both *LIPT1^-/-^*H460 and H157 cells, supporting a role for 2-HG-driven heterochromatin in fork impediment (Fig. 2B and D). Similarly, we previously reported that the constitutive heterochromatin mark H3K9me3 was elevated in *LIPT1^-/-^* cells compared with WT controls (*30*). Here, we found that the treatments with either α-KG or dimethyl-α-KG decreased H3K9me3 levels in *LIPT1^-/-^* H460 cells (Fig. S2A).

**Fig. 2.**
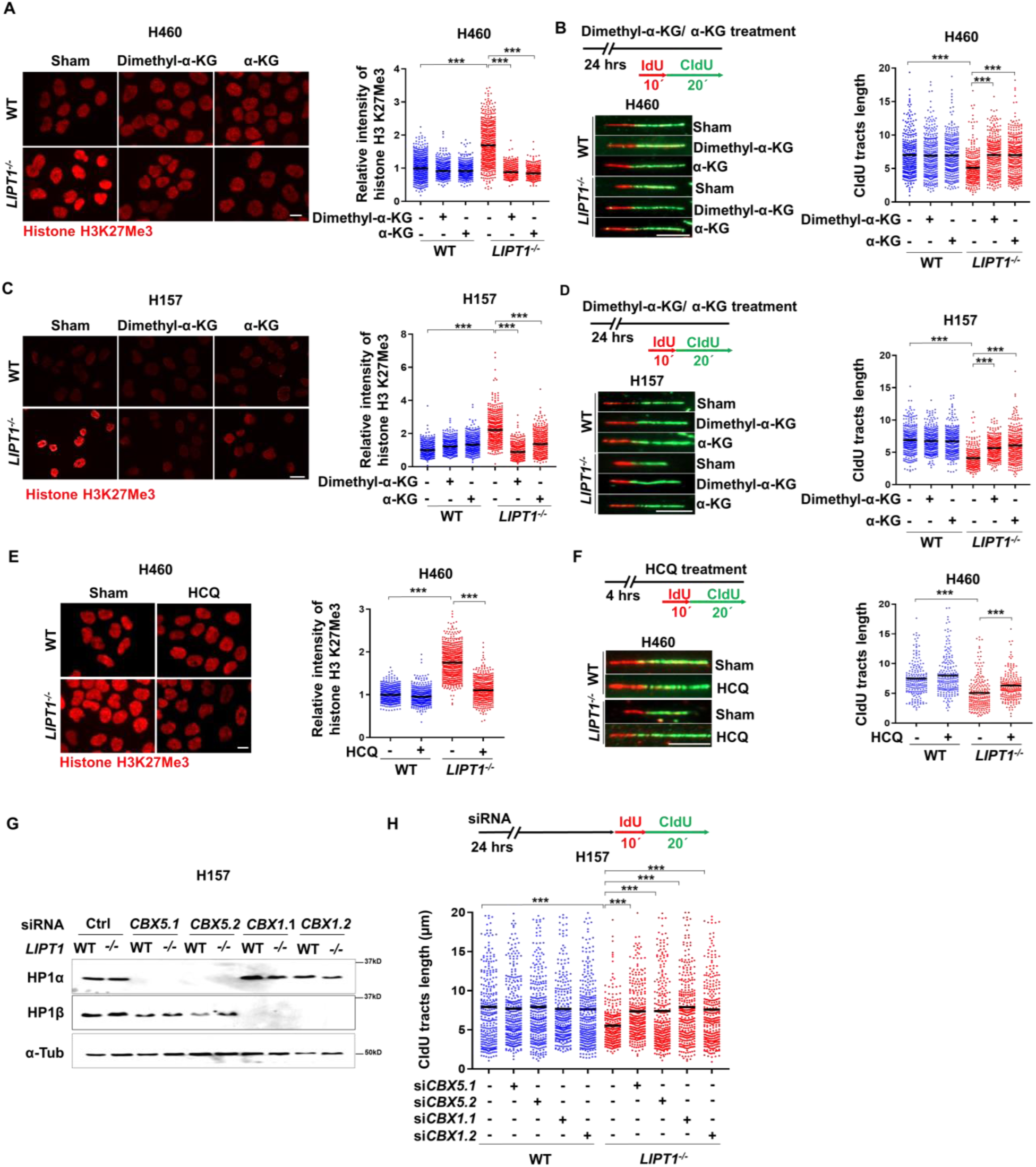
2-HG-mediated heterochromatin impedes replication fork in LIPT1-deficient cells. (**A**, **C, E**). Immunofluorescence of H3K27me3 in WT and *LIPT1^-/-^* H460 (A), H157 (C) cells pretreated with 1 mM dimethyl-α-KG or α-KG for 24 hours, or H460 cells ±10 µM hydroxychloroquine (HCQ) for 4 hours (E). Scale bar, 5 μm. (**B, D, F**). Top: DNA fiber assay of cells in A, C, E. Cells pretreated with dimethyl-α-KG/α-KG (B, D) or HCQ (F), and then pulse-labeled with IdU (10 min) and CldU (20 min). Representative fiber images (bottom) and quantification of CldU tract lengths (Right). Scale bar, 5 μm. (**G**) Immunoblot analysis of HP1α (encoded by *CBX5*) and HP1β (encoded by *CBX1*) in WT and *LIPT1^−/−^*H157 cells after siRNA knockdown. (**H**) DNA fiber assay of indicated cells labelled with IdU (10 min) and CIdU (20 min) at 24 hours after siRNA transfection. >600 cells per treatment (A, C, E), and >300 CIdU tracts lengths per condition were analyzed (B, D, F, H) from 3 independent experiments. Two-way ANOVA, \*\*\**P* < 0.001.

Hydroxychloroquine (HCQ), previously shown to reduce H3K27me3 and alleviate heterochromatin (*32-34*), also alleviated these replication defects. Specifically, pretreatment with 10 μM HCQ for 4 hours reduced H3K27me3 levels by immunofluorescence staining (Fig. 2E) and partially rescued replication fork progression (Fig. 2F) in *LIPT1^-/-^* H460 cells. Similarly, HCQ also reduced H3K27me3 by immunoblotting (Fig. S2B) and partially restored replication fork progression in *LIPT1^-/-^* HeLa cells (Fig. S2C). Heterochromatin protein 1 (HP1) is essential to promote chromatin compaction by directly binding to H3K9me3, which is further stabilized by H3K27me3 (*35*). We next performed DNA fiber assay after siRNA-mediated knockdown of HP1α (*CBX5*) and HP1β (*CBX1*) in *LIPT1^-/-^*H157 and HeLa cells. We found this intervention restored replication fork progression (Fig. 2G-H and S2D, E), supporting the role for heterochromatin in fork impediment.

### LIPT1-deficient cells exhibit replication fork stalling and replication stress

We next assessed whether the slowed replication observed in LIPT1 deficient cells leads to replication fork stalling and stress. DNA replication typically proceeds bidirectionally from a single origin, with both forks advancing symmetrically at similar speeds. However, fork asymmetry and stability can occur at the stalled forks (*36*). Consistent with replication stalling, we observed increased fork asymmetry measured by the ratio of longer fork over the shorter sister fork, in *LIPT1^-/-^* H460 and HeLa cells compared to their respective WT cells (Fig. 3A-B).

**Fig. 3.**
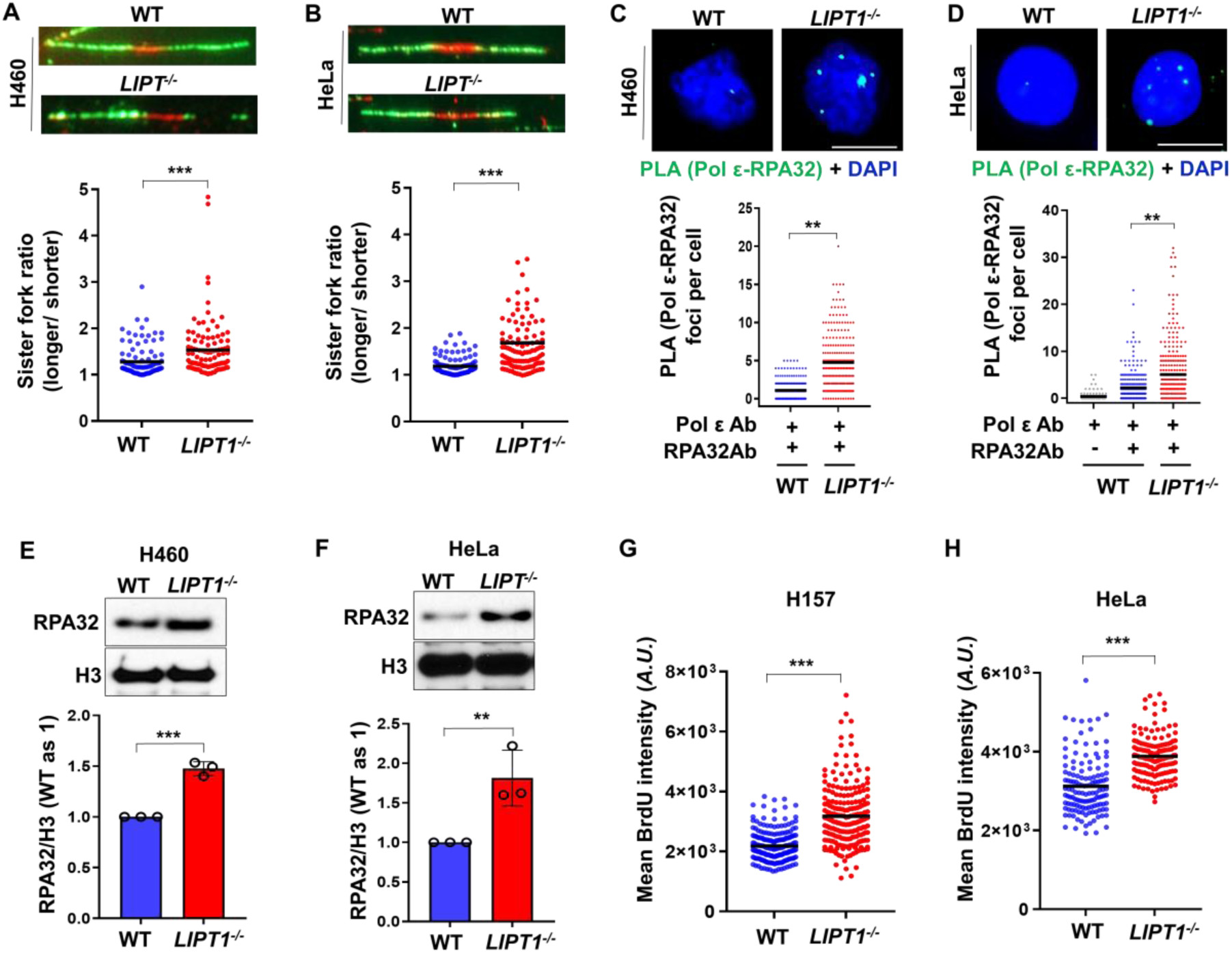
LIPT1-deficient cells exhibit replication fork stalling and replication stress. (**A-B**) Representative images and quantification of long arm over short arm of asymmetric sister replication forks in WT and *LIPT1^-/-^* H460 (A) and H157 (B) cells. (**C-D**) Representative images and quantification of PLA foci using antibodies against DNA polymerase ε (Pol ε) and RPA32 in WT and *LIPT1^-/-^* H460 (C) and HeLa (D) cells. Scale bars, 10 μm. >200 cells per condition from 3 independent experiments. (**E-F**) Immunoblot analysis of RPA32 in chromatin fractions of WT and *LIPT1^-/-^* H460 (E) and HeLa (F) cells. RPA32 levels were normalized to histone H3, and quantification from three independent experiments is shown as mean ± SD. (**G-H**) Immunofluorescence intensity of BrdU in PCNA-positive WT and *LIPT1^-/-^*H157 (G) and HeLa (H) cells under non-denaturing conditions. BrdU intensity was quantified in >100 PCNA-positive, S-phase cells per group. Wilcoxon rank-sum test was used, \*\**P* < 0.01 and \*\*\**P* < 0.001.

At the stalled fork, DNA polymerase pauses while helicase continues to unwind DNA, leading to the accumulation of single-stranded (ssDNA), which is rapidly coated by replication protein A (RPA) (*37, 38*). Thus, ssDNA in the vicinity of the stalled polymerase is frequently used as a marker of stalled fork and RS. Using proximity ligation assay (PLA) with antibodies against catalytic polymerase ε and RPA32, a key subunit of the RPA complex, we evaluated the presence of RPA-coated ssDNA in the vicinity of polymerase. We found that *LIPT1^-/-^* H460 and HeLa cells exhibit increased basal foci signal compared to their WT control cells (Fig.3C-D). In addition, we observed an increase of chromatin-bound RPA32 in *LIPT1^-/-^*H460 and HeLa cells compared to their WT control cells (Fig. 3E-F), suggesting increased recruitment of RPA32 to the exposed ssDNA.

To directly visualize ssDNA, we labeled genomic DNA with 5-bromo-2’-deoxyuridine (BrdU) over two consecutive cell cycles (48 hours), then washed the cells in BrdU-free medium and stained with BrdU antibodies. Under non-denaturing condition, BrdU antibodies only interact with the exposed BrdU in ssDNA but not dsDNA (*39*). We found that loss of LIPT1 increased BrdU labeling in both H157 and HeLa cells. (Fig. 3G-H), suggesting increased ssDNA. Together, these data suggest that LIPT1 loss leads to replication fork stalling and RS.

### LIPT1 loss induces PrimPol-generated ssDNA gaps

To further assess ssDNA formation, we performed a modified DNA fiber assay in which CIdU labeled DNA fibers were treated with or without S1 nuclease, an enzyme that specifically cleaves ssDNA(*40*). In the presence of ssDNA, CIdU tracts length is expected to shorten. S1 nuclease treatment markedly reduced CIdU tracts length in *LIPT1^-/-^* H157 (Fig. 4A, Fig. S3B), H460 (Fig. 4D) and HeLa (Fig. 4F, Fig. S3D) cells, but not in their respective WT controls. Notably, pretreatment with dimethyl-α-KG treatment or knockdown of HP1α via *CBX5*-targeting siRNA partially reduced the S1-induced CIdU tracts shortening in *LIPT1^-/-^*H157 cells (Fig. 4A), suggesting that 2-HG-driven heterochromatin contributes to ssDNA formation.

**Fig. 4.**
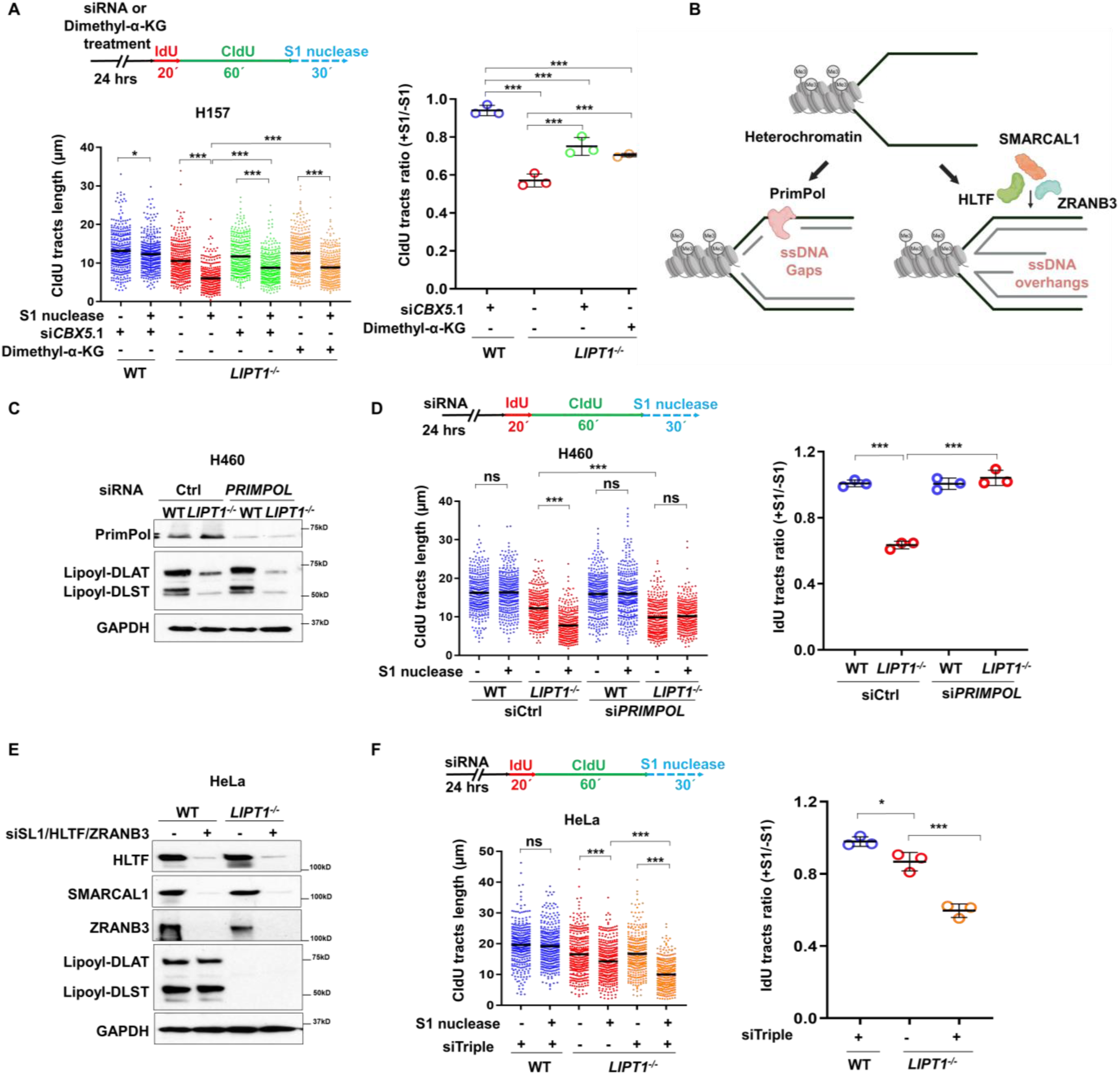
LIPT1 loss induces PrimPol-generated ssDNA gaps. (**A**) Top: DNA fiber assay of WT and *LIPT1^-/-^* H157 cells transfected with control or *CBX5* siRNAs, pretreated ± 1mM dimethyl-α-KG for 24 hours. Cells were sequential pulse labeled with IdU (10 min) and CldU (60 min), then treated ± S1 nuclease. Bottom: Quantification of CIdU tract lengths. Right: Ratio of S1-treated to untreated tract lengths. (**B**) Schematic model illustrating PrimPol-dependent repriming and replication fork reversal as competing mechanisms to resolve heterochromatin-induced replication stress. (**C**, **E**) Immunoblotting of PrimPol in WT and *LIPT1^-/-^* H460 cells (C), or HLTF, SMARCAL1, and ZRANB3 in WT and *LIPT1^-/-^* HeLa cells (E) after siRNAs transfection. (**D**, **F**) Top: DNA fiber assay of WT and *LIPT1^-/-^*H460 (D) and HeLa (F) cells 24 hours following siRNAs for 24 hours, labeled and treated as (A). Bottom: Quantification of CIdU tract lengths. Right: Ratio of S1-treated to untreated CIdU tract lengths. For A, D and F, >300 tracts/condition were analyzed across 3 independent experiments. Data are mean ± SD; significance by two-way ANOVA, \**P* < 0.05 and \*\*\**P* < 0.001.

At stalled forks, DDT mechanisms allow cells to bypass replication impediments and resume DNA synthesis, processes often generate ssDNA. Two major DDT pathways are commonly employed. PrimPol-mediated repriming restarts DNA synthesis downstream of blocking lesions, leaving behind post-replicative ssDNA gaps (Fig. 4B) (*41-43*). In contrast, the helicase-like enzymes SMARCAL1, HLTF, and ZRANB3 mediate the reversal of the stalled fork into a four-way structure. Although fork reversal does not directly generate ssDNA, it can produce ssDNA overhangs due to strand length asymmetry or nucleolytic degradation (Fig. 4B) (*44, 45*).

To determine whether the ssDNA gaps were generated through PrimPol repriming, we depleted PrimPol using siRNA and performed an S1 nuclease-based DNA fiber assay. PrimPol knockdown almost completely reversed the S1-induced CIdU tracts shortening in LIPT1-deficient cells (Fig. 4C-D, S3A-D)., indicating that PrimPol-mediated repriming generated ssDNA gaps LIPT1. To assess the contribution of fork reversal, we performed the S1 nuclease assay following simultaneous knockdown of *SMARCAL1*, *HLTF*, and *ZRANB*3 (*44, 45*). Interestingly, depletion of these factors did not alleviate but instead increased ssDNA gaps in LIPT1-deficient cells (Fig. 4E-F). Prior studies suggest that repriming and fork reversal compete at stressed forks, such that inhibition of one can enhance the other (*45-47*). Our data indicate that suppression of fork reversal may further upregulate PrimPol-mediated ssDNA gaps. Collectively, these findings identify PrimPol as the primary DDT mechanism in LIPT1 loss associated RS.

### PrimPol-generated ssDNA gaps require PARP1 for repair

Recent studies have shown that PARP plays a critical role in repairing PrimPol-mediated ssDNA gaps (*46, 48*). To examine this in LIPT1-deficient cells, we performed DNA fiber assays with sequential IdU and CldU labeling, followed by S1 nuclease digestion to monitor CIdU tract shortening in the presence or absence of PARP inhibitors (Fig. 5A). Tracts lengths were assessed 20 hours after labeling, a timeframe consistent with ssDNA gap repair or conversion to more complex double-strand breaks (DSBs) /chromosome lesions (*46*). In *LIPT1^-/-^* H460 and H157 cells, S1-induced tract shortening was largely resolved at 20 hours after labeling, whereas PARP inhibition completely abolished ssDNA gap resolution (Fig. 5B-C).

**Fig. 5.**
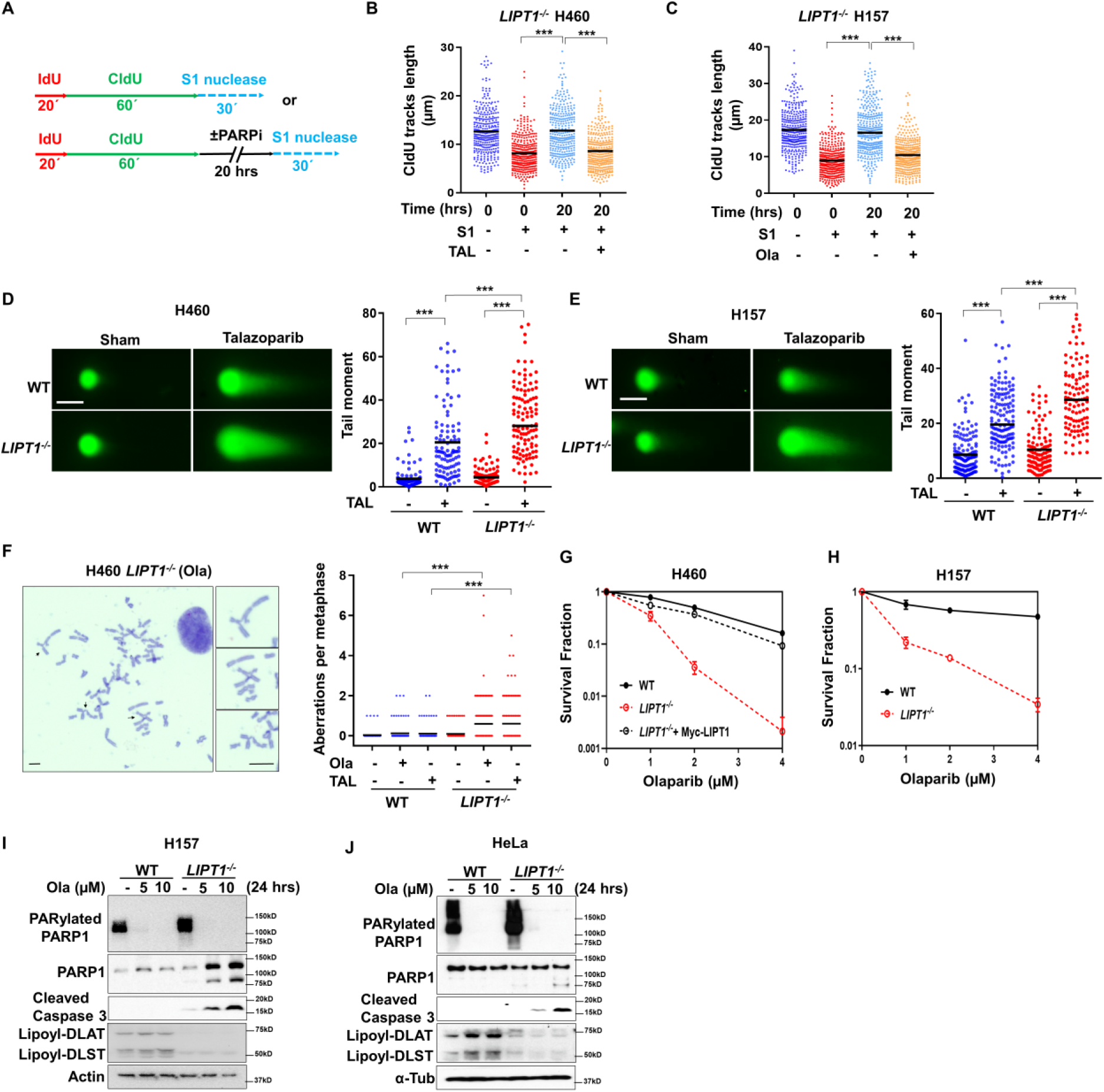
PrimPol-generated ssDNA gaps require PARP1 for repair. (**A**) DNA fiber assay for *LIPT1^-/-^*cells sequentially labeled with ldU and CIdU, then treated with S1 nuclease immediately or after 20 h recovery with or without PARP inhibitor (**B-C**). CldU tracts lengths in *LIPT1^-/-^* H460 (B, 100 nM talazoparib) or H157 (C, 10 µM olaparib) cells. >300 tracts per condition from 3 independent experiments were analyzed. Two-way ANOVA, \*\*\**P* < 0.001. (**D-E**) Neutral comet assays showing tail moment in WT and *LIPT1^−/−^*H460 (D) and H157 (E) cells after talazoparib (200 nM, 48hr). Scale bar, 50 μm. (**F**) Chromosome aberrations in WT and *LIPT1^-/-^* H460 cells after olaparib (10 µM, 48 hours); >100 cells; scale bar, 5 μm; Two-way ANOVA, \*\*\**P* < 0.001. (**G-H**) Clonogenic survival of WT and *LIPT1^-/-^* H460 (G) and H157 (H) cells after olaparib (10 µM, 48 hours); normalized to the DMSO-treated cells. (**I-J**). Immunoblot analysis of PAR, PARP1, cleaved Caspase-3, lipoyl-DLAT/DLST, and β-actin in WT and *LIPT1^−/−^* H157 (I) and HeLa (J) cells after olaparib (10 µM, 24hr).

Persistent ssDNA gaps can be converted into lethal DSBs or chromosome aberrations during subsequent DNA replication cycles (*49, 50*). To evaluate this consequence of PARP inhibition in context of LIPT1 deficiency, we performed neutral comet assays to assess DSBs formation in *LIPT1^-/-^* and WT cells treated with PARP inhibitors. Increased tail moment, reflecting elevated DSBs, was more pronounced in *LIPT1^-/-^* H460 cells after talazoparib (Fig. 5D) and in *LIPT1^-/-^* H157 after olaparib (Fig. 5E) compared with their WT counterparts. Cytogenetic analysis of metaphase spreads supported this effect, showing increased chromosome aberrations in *LIPT1^-/-^* H460 cells compared to WT controls after PARP inhibition (Fig. 5F).

Consistent with these findings, clonogenic assays showed that LIPT1 loss sensitized H460, H157, and HeLa cells to olaparib (Fig. 5G-H, S4A). Pretreatment with lipoylation inhibitor CPI-613 similarly enhanced sensitivity of these cells to olaprib, talazoparib or rucaparib (Fig. S4B-G). Western blots analysis revealed elevated basal PARylated PARP1 in *LIPT1^-/-^* H157 and HeLa cells relative to WT (Fig.5I-J), perhaps reflecting a compensatory response to repair the elevated basal ssDNA. Following 24-hour olaparib treatment, PARP1 PARylation is completely inhibited and accompanied by increased cleaved caspse-3 and cleaved PARP1 in *LIPT1^-/-^* H460 and H157 compared to WT controls, suggesting enhanced apoptosis.

Together, these results demonstrate that PrimPol-generated ssDNA gaps in LIPT1-deficient cancer cells require PARP1 for repair. Consequently, targeting lipoylation-deficient tumors with PARP inhibitors or other agents that exploit persistent ssDNA gaps represents a potential therapeutic strategy.

### PrimPol-generated nascent DNA undergoes MRE11-depedent resection

ssDNA gaps at the stalled replication forks, if not promptly repaired, can expose nascent DNA to nucleolytic degradation, resulting in the loss of newly synthesized DNA and genomic instability(*33, 51, 52*). To assess nascent DNA degradation, cells were sequentially labeled with IdU and CIdU, treated with 4mM hydroxyurea (HU) to induce RS and analyzed using DNA fiber assays. Degradation was indicated by a reduced CIdU/IdU ratio. (Fig. 6A). *LIPT1^-/-^*H460 and H157cells displayed markedly reduced CIdU/IdU ratios compared with WT controls (Fig. 6B, S5A), indicating that LIPT1 loss promotes nascent ssDNA degradation. PrimPol knockdown fully rescued this phenotype (Fig. 6B, S5A), whereas combined knockdown of the fork reversal factors HLTF, SMARCAL1, and ZRANB3 had no effect (Fig. 6C), confirming PrimPol as the source of the degraded nascent DNA.

**Fig. 6.**
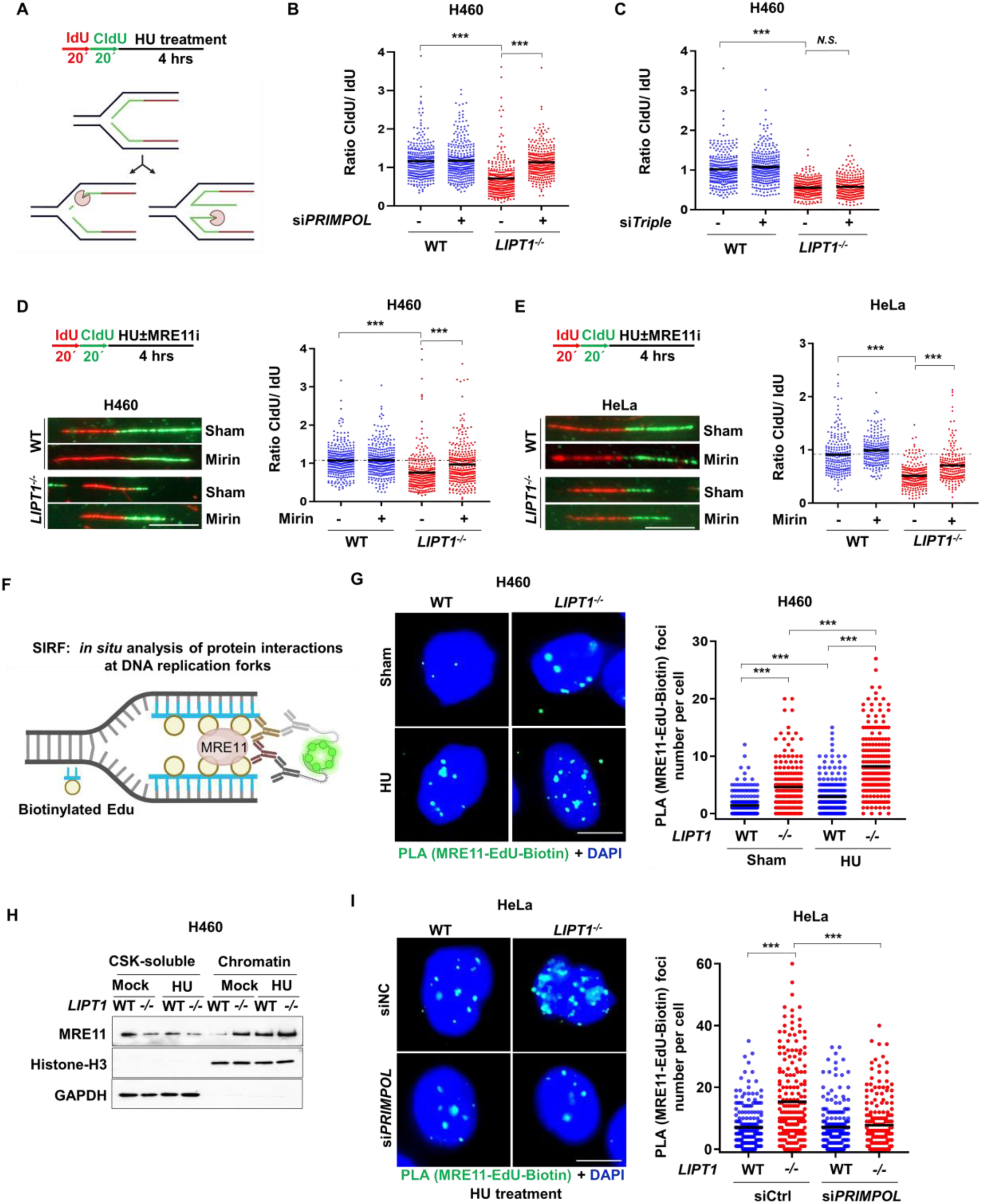
PrimPol-generated nascent DNA undergoes MRE11-depedent resection. (**A**) DNA fiber assay of cells sequentially labeled with IdU (20 min) and CldU (20 min), then treated with HU (4 mM, 4 h). Nascent DNA degradation from ssDNA (gaps and ends from reversed fork illustrated) was assessed by CIdU/IdU ratios. (**B**-**C**) CIdU/IdU ratios in WT and *LIPT1^-/-^* H460 cells after *PRIMPOL* (B) or *HLTF*/*SMARCAL1*/*ZRANB3* (C) knockdown; (**D-E**) DNA fiber degradation in WT and *LIPT1^-/-^* H460 (D) and H157 (E) cells ± Mirin (50 µM, 4 h). Representative fibers and quantification shown. Scale bar, 5 μm. (**F**) Schematic of SIRF assay: cells pulse-labeled with EdU (100 μM, 10 min), biotinylated via click chemistry, and analyzed by PLA with antibodies against biotin and MRE11; (**G**) Representative images and quantification of MRE11 by SIRF in WT and *LIPT1^-/-^*H460 cells. Scale bars, 10 μm. (**H**) Immunoblot of MRE11 in soluble (GAPDH marker) and chromatin-bound (histone H3 markers) fractions from WT and *LIPT1^-/-^* H460 cells ± HU (2 mM, 2 h). (**I**) Representative images and quantification of MRE11 by SIRF in WT and *LIPT1^-/-^* H460 cells after PRIMPOL knockdown. Scale bars, 10 μm. >300 DNA fiber (B-E) or >200 cells (G, I) from 3 independent experiments. Two-way ANOVA. \*\*\**P* < 0.001.

Previous studies have linked degradation of PrimPol-generated ssDNA to the exonuclease activity of Meiotic Recombination 11(MRE11) (*33, 51, 52*). To test the role of MRE11 in LIPT1-deficient cells, we performed fiber degradation assay with or without Mirin, a MRE11 inhibitor. Mirin treatment suppressed nascent DNA degradation in *LIPT1^-/-^* H460 and HeLa cells (Fig. 6D-E). We next examined MRE11 recruitment to nascent DNA using PLA-based, in <u>S</u>itu detection of Protein-DNA Interaction at <u>R</u>eplication <u>F</u>orks (SIRF) (*53*) (Fig. 6F). WT and *LIPT1^-/-^* H460 were cells pulse-labeled with EdU (100 μM, 10 min), biotinylated via click chemistry, and analyzed by PLA with antibodies against biotin and MRE11. *LIPT1^-/-^* cells exhibited increased MRE11: biotin-EdU PLA foci both at baseline and following HU, indicating enhanced MRE11 association with nascent DNA in *LIPT1^-/-^*cells (Fig. 6G, S5B). Chromatin fractionation confirmed elevated chromatin-bound MRE11 in LIPT1-deficient H460 and H157 cells under both basal and HU-treated conditions, with fraction purity verified by histone H3, and absence of GAPDH (Fig. 6G, S5E). Importantly, PrimPol knockdown markedly reduced MRE11 localization to EdU-labeled nascent DNA in *LIPT1^-/-^* cells, restoring it to WT levels (Fig. 6H).

Together, these results demonstrate that LIPT1 loss induces nascent DNA degradation. The degraded DNA originates from PrimPol-mediated repriming and is processed by MRE11-dependent resection.

### LIPT1 loss induces genomic instability

Both PrimPol-generated ssDNA gaps and degradation of nascent DNA pose a major threat to genomic stability (*46, 47, 54, 55*). Unresolved ssDNA gaps can persist into mitosis, leading to γH2AX accumulation at fragile chromosomal sites (*1*). Consistent with this, *LIPT1^-/-^* H460 and HeLa cells exhibited increased γH2AX foci in mitotic cells, identified by histone H3Ser10 phosphorylation and condensed chromosomes, compared to controls (Fig. 7A, 7C).

**Fig. 7.**
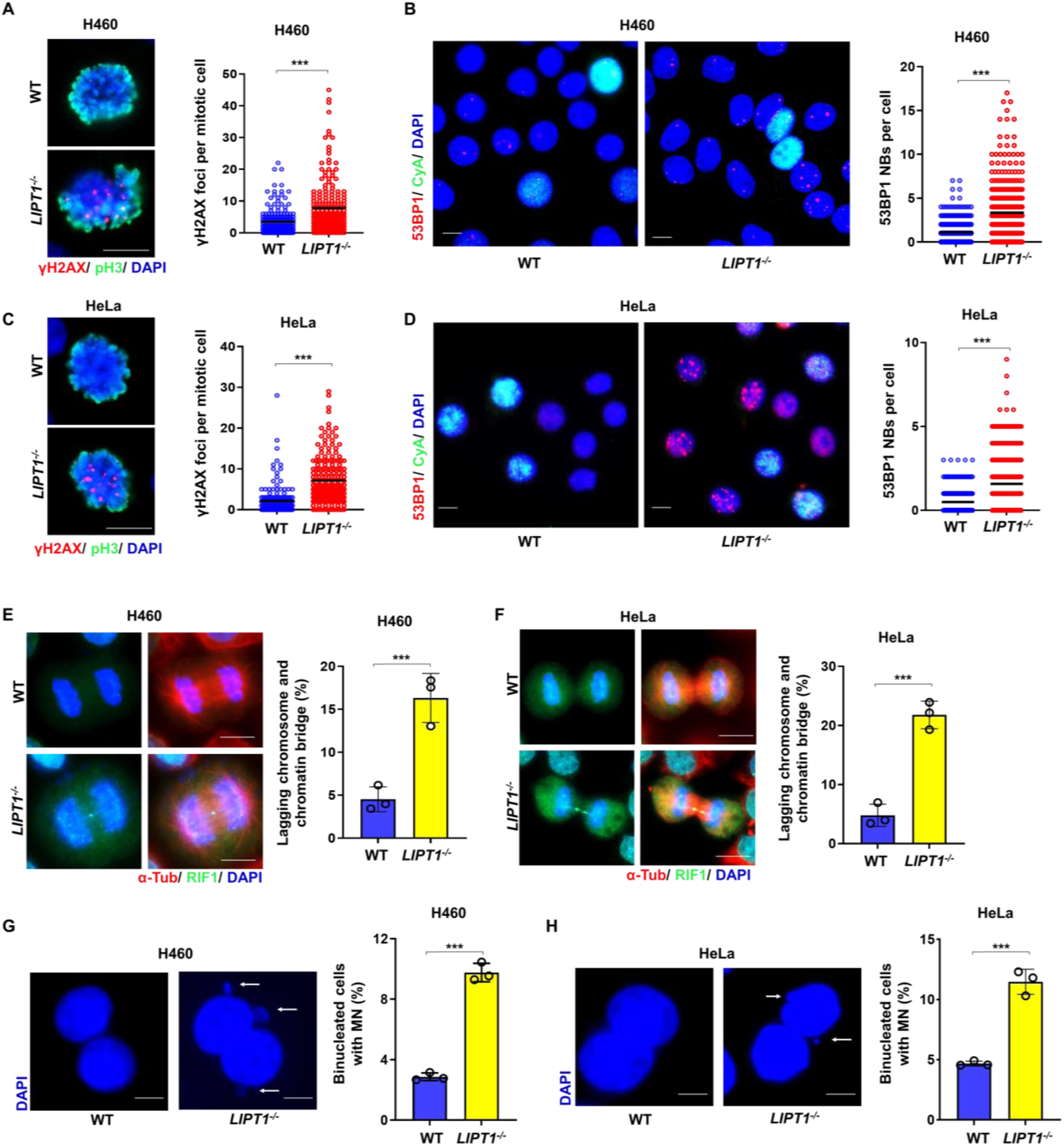
LIPT1 loss induces genomic instability. (**A**) Representative image and quantification of γH2AX foci in prometaphase of WT and *LIPT1^-/-^* H460 cells; scale bar, 10 μm; >100 prometaphase cells per group. (**B**) Representative image and quantification of 53BP1 nuclear bodies (NBs) in G1-phase nuclei of WT and *LIPT1^-/-^* H460 cells, identified by the absence of cycle A staining. Scale bar, 10 μm; >400 G1-phase cells per group. (**C-D**) γH2AX foci in prometaphase (C) and 53BP1 NBs in G1-phase nuclei (D) in WT and *LIPT1^-/-^* HeLa. (**E-F**) Immunofluorescent staining of WT and *LIPT1^-/-^* H460 (E) and HeLa (F) cells with anti-RIF1 (green) and anti-α-tubulin (red) antibodies. Representative images of anaphase cells with ultrafine chromatin bridges are shown in left panel. Right: percentage of anaphase cells with ultrafine chromatin bridges. (**G-H**) Representative image and quantification of micronuclei in binucleated WT and *LIPT1^-/-^*H460 (G) and HeLa cells (H) after cytochalasin B treatment (3 µg/mL, 40 hours) to block cytokinesis. Data are presented as mean ± SD from 3 independent experiments, with at least 100 anaphase cells (E-F) and 200 binucleated cells (G-H) analyzed per group. Wilcoxon rank-sum, \*\*\**P* < 0.001.

Under-replicated DNA can be transmitted into the next cell cycle, where it is bound by 53PB1 and sequestered into nuclear bodies (NBs) during G1, serving as a protective mechanism until repair occurs in S phase (*56, 57*). Quantification of 53BP1 NBs in cyclin A-negative (G1 phase) cells revealed a marked increase in *LIPT1^-/-^*H460 and HeLa cells compared with WT controls (Fig. 7B, 7D). Incomplete replication or persistent ssDNA can also form ultrafine anaphase bridges (UFBs), thread-like DNA connections between sister chromatids that impede proper chromosome segregation. Indeed, LIPT1-deficient cells displayed increased UFBs marked by Rap1-Interacting Factor 1(RIF1), a factor that localizes to UFBs and helps resolve under-replicated DNA (*58, 59*) (Fig. 7E-F). These lagging chromosomes during anaphase can be enveloped by the nuclear membrane, forming micronuclei. Accordingly, *LIPT1^-/-^* H460 and HeLa cells showed an increase in micronuclei formation compared to WT controls (Fig. 7G-H). Collectively, these results indicate that LIPT1 is critical for maintaining genomic integrity.

In summary, our data define a key function of LIPT1 in safeguarding DNA replication and genomic integrity (Fig.8). LIPT1 loss induces 2-HG–driven heterochromatin formation, which stalls replication fork progression. PrimPol-mediated repriming restarts DNA synthesis but generates ssDNA gaps, which are further expanded by MRE11-dependent degradation of nascent DNA. Repair of these ssDNA gaps by PARP1 enables cell survival despite genomic instability. PARP inhibition prevents repair and causes PARP trapping, resulting in persistent ssDNA gaps that are converted into lethal DSBs and cell death by apoptosis.

**Fig. 8.**
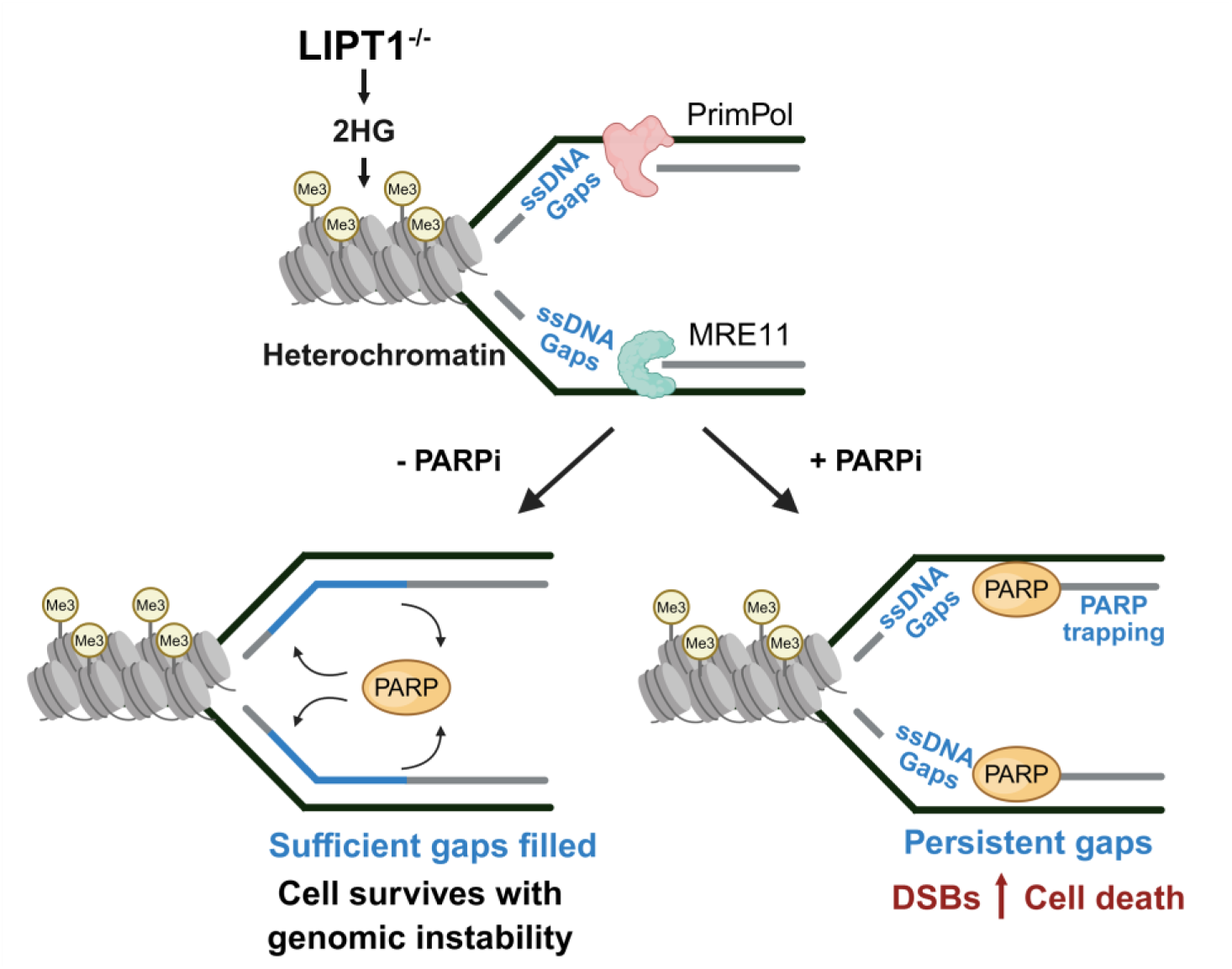
Schematic illustrating how LIPT1 deficiency promotes genomic instability and PARP inhibitor sensitivity. LIPT1 loss induces 2-HG–driven heterochromatin, which stalls replication fork progression. PrimPol-mediated repriming restarts DNA synthesis, generating ssDNA gaps that are further expanded by MRE11-dependent degradation of nascent DNA. Repair of ssDNA gaps by PARP1 enables cell survival despite genomic instability. PARP inhibition prevents repair, causing persistent ssDNA gaps that are converted into lethal DSBs.

## Discussion

DNA replication stress is a major threat to genomic integrity, arising from defects in DNA repair pathways or from physical impediments to fork progression, such as DNA adducts, secondary structures, or compact heterochromatin (*60, 61*). Our study reveals a critical role of lipoylation in preserving DNA replication and genome integrity. Together with prior literature, our findings support a multi-pronged mechanism by which lipoylation deficiency promotes genomic instability. First, loss of lipoylation leads to accumulation of 2-HG, which drives heterochromatin formation and imposes barriers to replication fork progression. Second, lipoylation-deficient cells depend on PrimPol-mediated repriming to restart stalled forks, generating ssDNA gaps that are further extended by MRE11. These gaps destabilize fork and predispose them to collapse (*33, 52, 62*), ultimately giving rise to anaphase bridge, 53BP1 nuclear bodies in subsequent replication cycles. When combined with PARP1 inhibition, they result in increased lethal DSBs. Third, aberrant chromatin remodeling impairs the recruitment of DNA repair machinery (*30, 31*), thereby compromising DNA damage repair signaling and the resolution of replication-associated lesions.

Oncogene-induced replication stress and genomic instability are well-established drivers of early tumorigenesis. For example, in non-small cell lung cancer (NSCLC), loss of BRG1 through *SMARCA4* mutation disrupts the SWI/SNF chromatin remodeling complex and promotes heterochromatin-associated replication stress in lung adenocarcinoma (*63, 64*), while activating KRAS mutation induces replication stress through H3K27me3- and HP1-dependent heterochromatin formation (*34*). Beyond tumor initiation, genomic instability also has prognostic relevance, as chromosomal instability in NSCLC correlates with increased recurrence and mortality (*65, 66*). In this context, our findings raise the possibility that lipoylation deficiency-induced replication stress and genomic stability may function as an oncogenic driver and prognostic marker. Supporting this, reduced expression of LIPT1 has been reported in several cancers including bladder and lung cancer, with lower LIPT1 levels correlating with worse prognosis (*27*).

At stalled fork, cells rely on DDT pathways to resume DNA synthesis, offering potential therapeutic opportunities. The two main DDT mechanisms are fork reversal and PrimPol-mediated repriming (*43, 48*). In LIPT1-deficient cancer cells, we observed a predominant reliance on PrimPol-mediated restart. The reason for this preference remains unclear. We previously reported that LIPT1 deficiency impairs TIP60-ATM-Rad51 signaling (*30*), an axis known to mediate fork reversal by recruiting SMARCAL1 to stalled forks (*51, 67*). Thus, LIPT1 deficiency may impair fork reversal to some extent, resulting in compensatory overactivation of PrimPol repriming and aggravation of ssDNA gap formation.

Although the resolution of ssDNA gaps is not fully understood, our data indicate a requirement for PARP1, rendering LIPT1-deficient cells sensitive to PARP inhibition. This is consistent with prior studies showing PAPR inhibitors induce PrimPol-generated ssDNA gaps in other cancer models of replication stress (*46, 50*). Inhibition of PARP1 leads to its retention on DNA/chromatin due to failure of PARylation, a well-characterized phenomenon termed PARP trapping. This is potentially more deleterious than persistent SSBs in the absence of PARP1, as forks colliding with trapped PARP1 can collapse and other repair proteins are excluded from accessing from the damage site.

LIPT1 deficiency therefore confers a “BRCAness” phenotype, characterized by homologous recombination defect, heightened replication stress, and PARP inhibitor sensitivity (*44, 68, 69*). This phenotype parallels that observed in Isocitrate dehydrogenase (IDH)-mutant gliomas, implicating elevated 2-HG as a potential mechanistic link. Supporting a synthetic lethal interaction between LIPT1 deficiency and PARP inhibition, a CRISPR screen using olaparib as selection pressure identified LIPT1 alongside with DNA repair genes such as RAD51, ATM, BRCA1 and BRCA2 (*70*). Notably, sgRNAs targeting IDH1/2 were not depleted, suggesting additional mechanisms beyond 2-HG accumulation contribute to the “BRCAness” of LIPT1-deficient cells. In particular, lipoylation has well-established roles in mitochondrial redox homeostasis and cuproptosis (*22*), both may influence genome stability through mechanisms yet to be elucidated. Although clinical trials of PARP inhibitor in IDH-mutant cancers have demonstrated limited efficacy (*71, 72*), further investigations into the contribution of LIPT1 deficiency to genomic instability, as well as synthetic combinations with agents targeting ssDNA gaps, are warranted.

Beyond cancer, lipoylation’s role in genome maintenance has implications for rare genetic diseases. Loss-of-function mutations in LIPT1 cause a severe inborn error of metabolism (IEM), often leading to fatal childhood crises marked by metabolic acidosis, encephalopathy, and multi-organ dysfunction (*23, 25*). While cancer risk in these patients cannot be assessed due to early mortality, our findings suggest that replication stress and genomic instability may contribute to disease pathogenesis, with genotoxic stressors acting as potential unrecognized triggers of fatal crises.

In summary, this study highlights lipoylation as a mechanistic link between mitochondrial metabolism, DNA replication, and genomic integrity. LIPT1 deficiency induces genomic instability that may act as both an oncogenic driver and a therapeutic vulnerability. These findings underscore the importance of further investigating lipoylation’s role in cancer progression and in the pathogenesis of lipoylation-related metabolic disease.

## Material and Methods

### Cell and cell culture

The human NSCLC cell lines H460 and H157 were obtained from J. Minna (Hamon Center Collection, UT Southwestern Medical Center. TX, USA) and cultured in RPMI 1640 medium (HyClone, Cytiva, Marlborough, MA) supplemented with 5% fetal bovine serum (FBS; Gemini Bio-Products, West Sacramento, CA), 2 mM glutamine, and 1% penicillin-streptomycin (P/S; Gibco, Thermo Fisher Scientific, Waltham, MA). The human cervical cancer cell line HeLa was obtained from the American Type Culture Collection and maintained in Dulbecco’s modified Eagle’s medium (DMEM) supplemented with 10% FBS and 1% P/S. All cells were maintained in a humidified incubator at 37°C with 5% CO₂ and used within 15 passages after thawing from original stocks. Mycoplasma contamination was routinely monitored using a PCR detection kit (Bulldog Bio, Portsmouth, NH). All cell lines were authenticated by DNA fingerprinting and confirmed to be mycoplasma-free.

### Generation of *LIPT1* knockout and Myc-LIPT1 expressing cell lines

*LIPT1^-/-^* H460, H157, and HeLa cells were generated using CRISPR-Cas9 as described previously(*30*). sgRNA targeting *LIPT1* (TGG TAG CCT GCA CAT CCA GC) and a non-targeting control (TTC TTA GAA GTT GCT CCA C) were cloned into the PX458 (Addgene #48138) and lentiCRISPR v2-Blast (Addgene #52962) vectors. H460 cells were transfected with control or *LIPT1* gRNA-pX458 constructs using Lipofectamine 3000 transfection reagent (Thermo Fisher Scientific, Waltham, MA), and GFP-positive cells were sorted by the BD FACSAria III Cell Sorter (BD biosciences, Franklin Lakes, NJ) for clonal expansion. H157 and HeLa cells were transduced with *LIPT1* gRNA-lentiCRISPR v2 particles produced in HEK293T cells, selected with 10 μg/ml blasticidin, and single-cell clones were established. *LIPT1* knockout clones were cultured in medium supplemented with 100 μg/ml uridine (MilliporeSigma, Burlington, MA) and 1 mM sodium pyruvate (MilliporeSigma) and validated by loss of DLAT and DLST lipoylation. For rescue, *LIPT1^-/-^* H460 cells were transduced with pLenti-EF1a-C-Myc-DDK-IRES-Puro-LIPT1-WT(*21*), selected with puromycin (1 μg/ml), and re-expression confirmed by anti-myc immunoblotting (#2276, Cell Signaling Technology, Danvers, MA).

### RNA interference

Transient knockdown was performed using Lipofectamine 3000 (L3000150, Invitrogen, Thermo Fisher Scientific, Waltham, MA) according to the manufacturer’s instructions. Briefly, 250 μL Opti-MEM (31985062, Gibco, Thermo Fisher Scientific, Waltham, MA) was mixed with 4 μL Lipofectamine 3000 and 10nM of indicated siRNA (sequences provided in Table S1). Cells were harvested for downstream assays 24 h post-transfection unless otherwise noted.

### Cell proliferation and clonogenic survival assays

For the cell viability assays, 1,000 cells were seeded in 96-well plates, and cell numbers were quantified at indicated time points using a Cytation 5 Multi-Mode Reader (BioTek, Winooski, VT, USA). For clonogenic survival assays, cells were seeded in 60-mm dishes (Greiner Bio-One, Kremsmünster, Austria) at appropriate densities and cultured for 7-10 days. For PARP inhibitors treatment, cells were exposed to PARP inhibitors at indicated concentration for 7-10 days. To assess combinational effects, cells were co-treated with CPI-613 and PARP inhibitors for 48 h, followed by continued exposure to PARP inhibitors alone for the remaining duration of the assay. Colonies were fixed and stained with crystal violet, and those containing more than 50 cells were counted as surviving colonies. Plating efficiency was normalized to either untreated controls or CPI-613-only controls to generate survival curves.

### Click-iT EdU staining assay

DNA synthesis was assessed using the click-iT^TM^ EdU cell proliferation kit (C10338, Thermo Fisher Scientific, Waltham, MA). Briefly, *LIPT1^-/-^* and WT H460 and H157 cells were seeded on coverslips in 6-well plates, incubated with 10 µM EdU for 30 min. Cells were fixed with 4% paraformaldehyde (PFA) in phosphate-buffered saline (PBS) for 20 min, permeabilized with 0.5% Triton X-100 in PBS for 15 min, and then incubated for 30 min in the Click-iT reaction cocktail (1×Click-iT reaction buffer, CuSO_4_, Alexa Fluor azide, and ×reaction buffer additive) protected from light. Nuclei were stained with 4′,6-diamidino-2-phenylindole (DAPI) for 10 min, and coverslips were mounted using antifade mounting medium (Vector Laboratories, Burlingame, CA) and sealed with nail polish. Images were acquired using an Axio Imager M2 fluorescence microscope (Carl Zeiss, Thornwood, NY, USA) and analyzed with AxioVision SE64 Rel.4.8 software (Carl Zeiss, Thornwood, NY, USA). Fluorescence intensity was quantified in ≥100 cells per condition across three independent experiments.

### Immunoblot analysis

Whole-cell lysates were prepared using lysis buffer (50 mM Tris, pH 7.5, 0.2 M NaCl, 1% Tween-20, 1% NP-40, 1 mM sodium orthovanadate, 2 mM β-glycerophosphate, and protease inhibitors). After centrifugation to clear the lysate, the protein concentration of each sample was measured using the Pierce BCA Protein Assay kit (23227, Thermo Fisher Scientific, Waltham, MA). SDS–polyacrylamide gel electrophoresis was performed using equal amount of protein, and the separated proteins were transferred onto the nitrocellulose membrane (Bio-Rad, Hercules, CA). The membranes were blocked using non-fat milk in tris-buffered saline containing 0.1% Tween-20 (Bioworld, Dublin, OH) and then incubated with primary antibodies overnight at 4°C. The nitrocellulose-bound primary antibodies were washed and incubated with horseradish peroxidase–linked secondary antibodies (Cell Signaling Technology, Danvers, MA). The immunoblots were reacted using Pierce enhanced chemiluminescent Western Blotting Substrate (Thermo Fisher Scientific, Waltham, MA), exposed to x-ray films, and then developed using a Protec x-ray film processor. Primary antibodies are listed in Table S2.

### Immunofluorescence staining

For immunofluorescence staining of H3K9me3/27, γH2AX, pH3Ser10, 53BP1, cyclin A, 5-bromo-2′-deoxyuridine (BrdU), RIF1, and α-Tubulin, cells were seeded on coverslips in six-well plates and cultured overnight. Cells were fixed with 4% paraformaldehyde (PFA) for 20 min at room temperature, permeabilized with 0.5% Triton X-100 in phosphate-buffered saline (PBS) for 15 min and blocked with 5% bovine serum albumin (BSA) in PBS for 30 min at room temperature. Primary antibodies (**Table S2**) were diluted in 1% BSA in PBS and incubated with cells overnight at 4 °C. After PBS washes, cells were incubated with fluorescent secondary antibodies for 1 hour at room temperature. Nuclei were counterstained with DAPI for 10 min. Coverslips were then mounted onto glass slides using antifade mounting medium (Vector Laboratories, Burlingame, CA) and sealed with nail polish. Images were captured using Axio Imager M2 fluorescence microscope (Carl Zeiss, Thornwood, NY, USA) and analyzed using AxioVision SE64 Rel.4.8 software (Carl Zeiss, Thornwood, NY, USA). For H3K9me3/27 staining, >600 cells per condition were quantified across three biological replicates. For BrdU incorporation to detect ssDNA, cells were pre-labeled with BrdU for 48 h, and BrdU fluorescence intensity was quantified in >200 cells per condition across two replicates. For prometaphase γH2AX staining, the number of γH2AX foci in each prometaphase cell was counted. For G1-phase 53BP1 nuclear body (NB) analysis, the number of 53BP1 NBs was counted in each cyclin A-negative cell. Each condition was evaluated in three biological replicates, with more than 200 prometaphase cells analyzed per condition. For ultra-fine chromatin bridge analysis, the percentage of anaphase cells containing RIF1-stained chromatin bridges was quantified in three biological replicates, with more than 100 anaphase cells counted per replicate.

### DNA fiber assay

Cells were sequentially labeled with 40 μM iododeoxyuridine (IdU) and 100 μM chlorodeoxyuridine (CldU) as indicated in the figure legends. Cells were harvested in cold PBS (1 × 10⁶/ml), and 3 μl of suspension was mixed with 8 μl spreading buffer (0.5% SDS, 200 mM Tris-HCl pH 7.4, 50 mM EDTA) and spread onto tilted glass slides. DNA fibers were fixed (methanol:acetic acid, 3:1, 10 min), denatured (2.5 N HCl, 60 min), and blocked (3% BSA/0.05% Tween-20, 30 min, 37 °C). CldU and IdU were detected using rat anti-BrdU (1:100; OBT0030, Accurate Chemical & Scientific Corporation, Carle Place, NY) and mouse anti-BrdU (1:50; 347580, BD Biosciences, San Diego, CA) for 1 hour, followed by goat anti-rat Alexa Fluor 488 (1:100; A-11006, Thermo Fisher Scientific, Waltham, MA) and goat anti-mouse Alexa Fluor 568 (1:100; A-11004, Thermo Fisher Scientific, Waltham, MA) for 1 hour at 37 °C. Slides were mounted with antifade medium (Vector Laboratories, Burlingame, CA), sealed, and imaged on an Axio Imager M2 fluorescence microscope (Carl Zeiss, Thornwood, NY, USA). Images were processed with AxioVision SE64 Rel.4.8 software (Carl Zeiss). S1 nuclease DNA fiber assays was performed as previously described(*73*). Following CIdU labeling, cells were permeabilized in CSK100 buffer (100 mM NaCl, 10 mM MOPS pH 7, 3 mM MgCl₂, 300 mM sucrose, 0.5% Triton X-100) for 10 min at room temperature and rinsed with PBS. Cells were then washed in S1 nuclease buffer pH 4.6 (30 mM NaAc, 10 mM ZnAc, 5% glycerol, 50 mM NaCl) and incubated with S1 nuclease (20 U/mL) for 30 min at 37 °C. After washing with PBS/0.1% BSA, nuclei were collected and processed for staining as the above DNA fiber assay.

### Neutral and Alkaline comet assay

The comet assays were conducted following the manufacturer’s instructions (#4250-050-K, R&D Systems, Minneapolis, MN). *LIPT1^-/-^* and WT H460 and H157 cells were harvested at 48 hours after talazoparib treatment, and 500 cells per condition were embedded in low-melting point CometAssay LMAgarose (#4250-050-02, R&D Systems). The agarose-cell suspension was spread onto microscope slides to form a thin layer and allowed to solidify. Slides for neutral comet assays were then immersed in ice-cold lysis buffer for 1 hour at 4°C, followed by a wash and immersion in 1× tris-borate-EDTA (TBE) buffer for 15 min. Slides for alkaline comet assays were incubated in ice-cold lysis buffer overnight at 4°C and then placed in freshly prepared alkaline unwinding solution (300 mM NaOH, 1 mM EDTA, pH>13) for 1 hour at 4°C. Electrophoresis was performed in a horizontal gel electrophoresis apparatus with freshly prepared 1× TBE for the neutral comet assay or alkaline electrophoresis solution (300 mM NaOH, 1 mM EDTA, pH >13) for the alkaline comet assay at 23 volts for 30 min at 4°C. The slides were then washed sequentially with dH₂O and 70% ethanol, air-dried at 37°C, and stained with SYBR Gold Nucleic Acid Gel Stain (Thermo Fisher Scientific, Waltham, MA) in Tris-EDTA buffer. DNA fragments were visualized using a Keyence BZ-X700 All-in-one Fluorescence microscope (Keyence, Osaka, Japan). Quantification of DNA damage was achieved by measuring parameters such as tail moment, defined as the product of the tail length and the percentage of DNA in the comet tail (Tail moment = tail length × % of DNA in the tail). This analysis was performed using OpenComet (v1.3)(*74*). Each experimental condition was assessed using two to three biological replicates with more than 100 cells analyzed per replicate.

### Cytokinesis-block micronucleus (CBMN) assay

Cells were treated with 3 μg/mL cytochalasin B (250233, MilliporeSigma, Burlington, MA) to block cytokinesis. After 24-48 h of incubation to allow nuclear division without cytokinesis, cells were harvested and fixed in methanol:acetic acid (3:1). Nuclei and micronuclei were visualized by DAPI staining and binucleated cells were analyzed microscopically. Micronuclei were quantified in ≥ 500 binucleated cells per condition, in three independent experiments.

### Metaphase chromosome spread assays

Chromosome spread assays were performed as previously described (*75*). WT and *LIPT1^-/-^* H460 cells were treated with 10 µM olaparib or 200 nM talazoparib for 48 h, harvested and incubated in pre-warmed 0.075 M KCl at 37 °C for 15 minutes to induce hypotonic swelling. Pre-fixation was performed by adding 1 ml of ice-cold Carnoy’s fixative (methanol:acetic acid, 3:1) directly to the suspension for 5 minutes on ice. Cells were then fixed in Carnoy’s solution overnight, followed by two additional 20-minute on ice. Fixed cells were dropped onto pre-warmed glass slides, air-dried overnight, stained with 5% Giemsa for 15 minutes, rinsed with water and mounted. Chromosome spreads were imaged using an Axio Imager M2 fluorescence microscope (Carl Zeiss, Thornwood, NY, USA) and analyzed with AxioVision SE64 Rel.4.8 software. Chromosomal aberrations, including breaks, deletions, and chromatid interchanges, were quantified as described(*76*). Each condition was evaluated in three biological replicates with ≥50 cells analyzed per replicate.

### Chromatin fractions isolation (CSK extraction method)

Cells were lysed in CSK-100 buffer (100 mM NaCl, 300 mM sucrose, 3 mM MgCl2, 10 mM PIPES pH 6.8, 1 mM EGTA, 0.2% Triton X-100) containing protease and phosphatase inhibitors at 4 °C for 15 min. Chromatin-associated proteins were released from the pellets by treatment with lysis buffer (50 mM HEPES pH 7.5, 50 mM NaCl, 0.05% SDS, 2 mM MgCl2, 10% Glycerol, 0.1% Triton X-100, 10 units of Benzonase Nuclease) containing protease inhibitors at 4 °C overnight. The supernatants were separated by SDS-PAGE and detected by immunoblotting with the indicated antibodies.

### Subcellular fractionation

Chromatin-bound RPA32 was examined using the Subcellular Protein Fractionation Kit (78840, Thermo Fisher Scientific, Waltham, MA). Cells were harvested and lysed using cytoplasmic extraction buffer and membrane extraction buffer. The pellets were first incubated with nuclear extraction buffer (NEB) to obtain the nuclear-soluble fraction, followed by treatment with NEB containing micrococcal nuclease and CaCl_2_ to extract chromatin-bound proteins. The protein lysates obtained from these fractions were analyzed using gel electrophoresis and Western blotting analysis.

### In situ Proximity ligation assay (PLA)

In situ PLA was performed using Duolink PLA reagents (DUO96020, MilliporeSigma, Burlington, MA) following the manufacturer’s instructions. Cells were extracted with CSK buffer (0.2% Triton X-100, 20 mM HEPES-KOH pH 7.9, 100 mM NaCl, 3 mM MgCl_2_, 300 mM sucrose, 1 mM EGTA) for 5 min, fixed with 4% paraformaldehyde (PFA) for 20 min at room temperature, and permeabilized with 0.5% Triton X-100 in phosphate-buffered saline (PBS) for 15 min. The cells were then treated with RNaseA (10 µg/ml) for 1 hour at 37 °C, washed, and blocked with Duolink in situ blocking solution for 1 hour. Cells were incubated overnight at 4°C with antibodies for Pol ε catalytic subunit (NBP2-55332, Novus Biologicals, Centennial, CO) and RPA32 (sc-56770, Santa Cruz Biotechnology, Dallas, TX) (1:500 diluted in Duolink in situ antibody diluent), then incubated with oligonucleotides-conjugates secondary antibodies (PLA probe anti-rabbit PLUS and anti-mouse MINUS), followed by ligation and amplification with a fluorophore-labeled oligonucleotide probe (excitation = 490 nm, emission = 520 nm). Images were captured with a fluorescence microscope (Axio Imager M2, Carl Zeiss, Thornwood, NY, USA) and were recorded using AxioVision SE64 Rel.4.8 software (Carl Zeiss, Thornwood, NY, USA).

### In situ analysis of protein interactions at DNA replication forks (SIRF) assay

SIRF assay was performed as previously described(*53*), with the following specifications. Cells were incubated with 100 µM EdU for 10 min and treated with 4 mM HU for 2 hours. Cells were sequentially pre-extracted at room temperature for 5 min each with CSK buffer (100 mM NaCl, 300 mM sucrose, 3 mM MgCl₂, 10 mM PIPES pH 6.8, EDTA-free protease inhibitor) and CSK-T (CSK + 0.5% Triton X-100). Cells were fixed with 4% PFA/PBS for 20 min and permeabilized with 0.25% Triton X-100 for 15 min at room temperature. Cells were washed 3 times for 5 min with PBS and incubated with fresh prepared Click-iT reaction buffer (2 mM copper sulfate, 10 µM biotin-azide, 100 mM sodium ascorbate in PBS) for 1 hour at room temperature in a humid chamber. Subsequently, cells were washed 2 times with PBS for 5 min and incubated in blocking buffer (10% goat serum and 0.1% Triton X-100) for 1 hour at room temperature. Anti-MRE11 (1:200; #4847, Cell Signaling Technology, Danvers, MA) and anti-Biotin (1:200; 03-3700, Thermo Fisher Scientific, Waltham, MA) antibodies were diluted in the blocking buffer and incubated overnight at 4°C. Cells were then processed for PLA staining as described above.

### Statistical analysis

Statistical analysis was performed using the Wilcoxon rank-sum test for two-group comparisons and two-way ANOVA for comparisons among more than two groups. All analyses were conducted with GraphPad Prism, with statistical significance defined as \**P* < 0.05; \*\**P* < 0.01 and \*\*\**P* < 0.001 were also reported where applicable.

## Acknowledgements

We thank Dr. Huiming Chen, Dr. Peter Ly for helpful discussions. Y.Z. is supported by Lung SPORE Career Enhancement Award, Distinguished Research Award from the President’s Research Council, Institutional Research Grant (IRG-21-142-16) from American Cancer Society, Cancer Center Support Grant (P30CA142543) from UT Southwestern and the Simmons Cancer Center, awards from the National Center for Advancing Translational Sciences of the National Institutes of Health (KL2TR003981 and CTSA-PP-YR1-D-009), and Startup Award from UT Southwestern Department of Radiation Oncology and Disease-oriented clinical scholar award (DOCS). A.J.D is supported by National Institutes of Health (R01CA276058 & R01CA29290), Department of Energy (DE-SC0025578), and National Aeronautics and Space Administration (20-20HHCSR_2-0033).

