## Supplementary material for "LIPT1 loss confers replication stress and PARP inhibitor sensitivity through PrimPol-mediated ssDNA gaps": Supplemental information.pdf

### Supplemental Figure S1

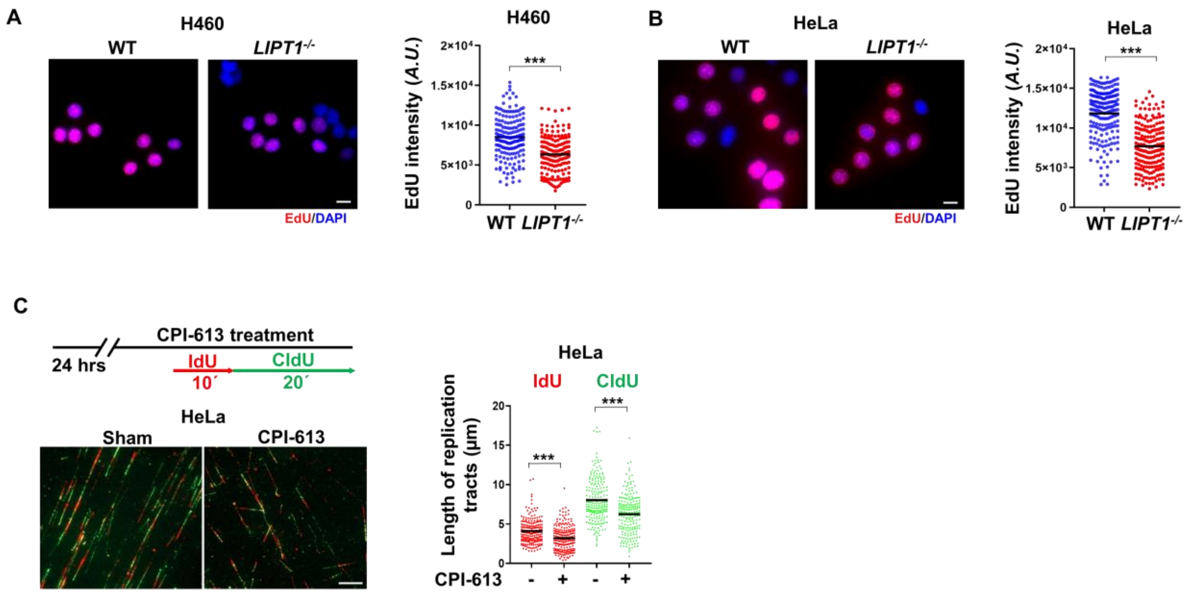


**Fig. S1. LIPT1 loss decreases DNA synthesis and replication.** WT and *LIPT1*$^{-/-}$ H460 (**A**) and HeLa cells (**B**) were pulse-labeled with 50 μM EdU for 30 min and EdU labeling was assessed via microscopy. Data are presented as EdU densities in nuclei (n > 100). (**C**) Representative images and quantification of IdU and CldU tracts lengths from the DNA fiber assay. HeLa cells were pretreated with DMSO or 100 μM CPI-613 for 24 hours, followed by sequential pulse-labeling with IdU (10 min) and CldU (20 min). The lengths of IdU (red) and CldU (green) tracts were measured from >200 ongoing DNA replication tracts across 3 independent experiments. Wilcoxon rank-sum test was used for comparisons between two groups, while two-way ANOVA test was applied for comparisons among more than two independent groups. Statistical significance is indicated as, $^{***}P < 0.001$.

### Supplemental Figure S2

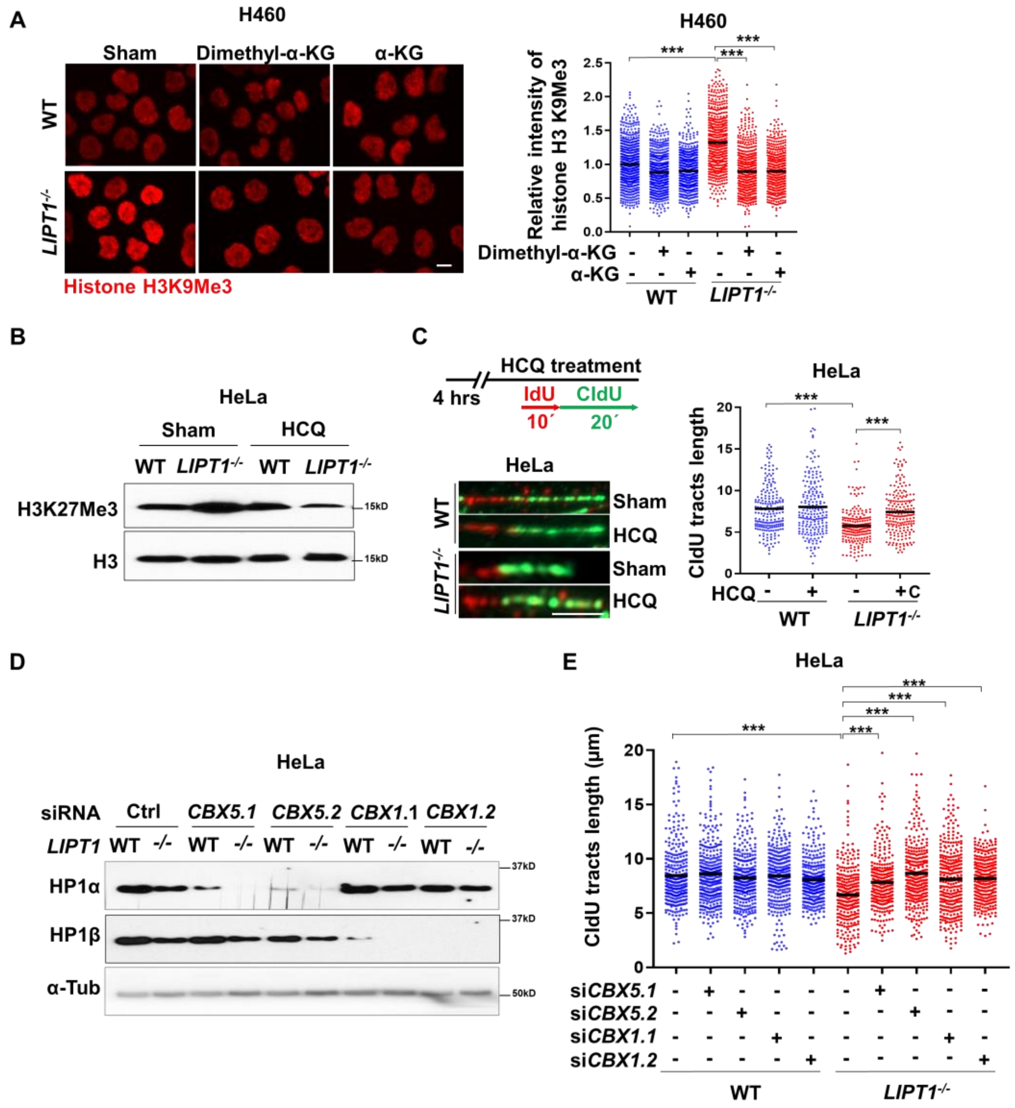


**Fig. S2.** (**A**) Immunofluorescence of H3K9me3 in WT and *LIPT1*$^{-/-}$ H460 cells pretreated with 1mM dimethyl-α-KG or α-KG for 24 hours. Nuclei were stained with Hoechst 33342. Scale bar, 5 μm. >600 cells per treatment from 3 independent experiments. Two-way ANOVA. ***$P < 0.001$. (**B**). Immunoblot analysis of H3K27me3 in WT and *LIPT1*$^{-/-}$ HeLa cells ±10 μM HCQ for 4 hours. Histone H3 was used as a loading control. (**C**) Top: DNA fiber analysis of WT and *LIPT1*$^{-/-}$ HeLa pretreated with10 μM HCQ for 4 hours, and then labelled with IdU (10 min) and CIdU (20 min). Representative fiber images (Bottom) and quantification of CldU tract lengths (Right). Scale bar, 5 μm. >300 ongoing DNA replication tracts from 3 independent experiments. Two-way ANOVA, $^{***}P < 0.001$. (**D**) Immunoblot analysis of HP1α (encoded by *CBX5*) and HP1β (encoded by *CBX1*) in WT and *LIPT1*$^{-/-}$ HeLa cells after siRNA knockdown. (**E**) DNA fiber assay of indicated cells labelled with IdU (10 min) and CIdU (20 min) at 24hr after siRNA transfection. >600 cells per treatment (A), and >300 CIdU tracts lengths per condition were analyzed (C, E) from 3 independent experiments. Two-way ANOVA, $^{***}P < 0.001$.

### Supplemental Figure S3

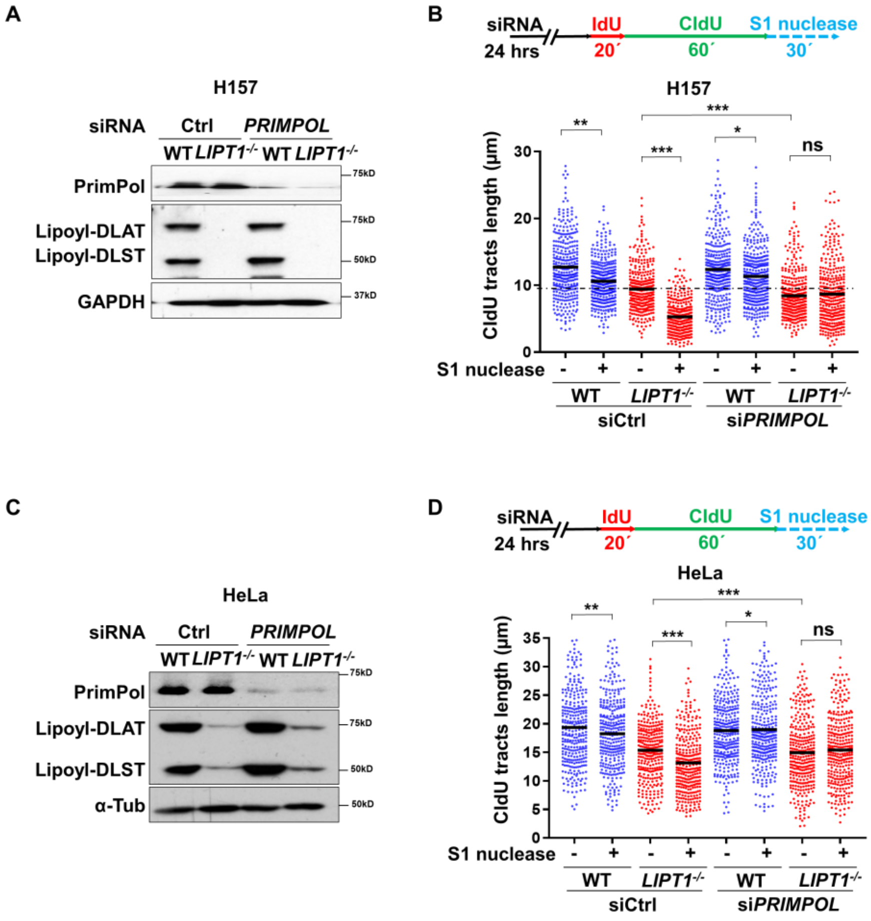


**Fig S3.** (**A**) Immunoblot analysis of PrimPol in WT and *LIPT1*$^{-/-}$ H157 cells following transfection with control or *PRIMPOL* siRNAs. (**B**) DNA fiber assay of WT and *LIPT1*$^{-/-}$ H157 cells transfected with control or *PRIMPOL* siRNAs for 24h, followed by IdU (20min) and CIdU (60min), then treated with or without S1 nuclease. (**C**) Immunoblot analysis of PrimPol in WT and *LIPT1*$^{-/-}$ HeLa cells following transfection with control or *PRIMPOL* siRNAs. (**D**) DNA fiber assay. WT and *LIPT1*$^{-/-}$ HeLa cells were treated the same as in B. *Bottom left*: Quantification of CldU tract lengths. *Right*: Ratio of S1-treated to untreated CldU tract lengths. For all fiber assays (A, D and F), >300 tracts per condition were analyzed across 3 independent experiments. Data are presented as mean ± SD. Statistical significance was assessed using two-way ANOVA, $^{*}P < 0.05$, $^{**}P < 0.01$, $^{***}P < 0.001$.

### Supplemental Figure S4

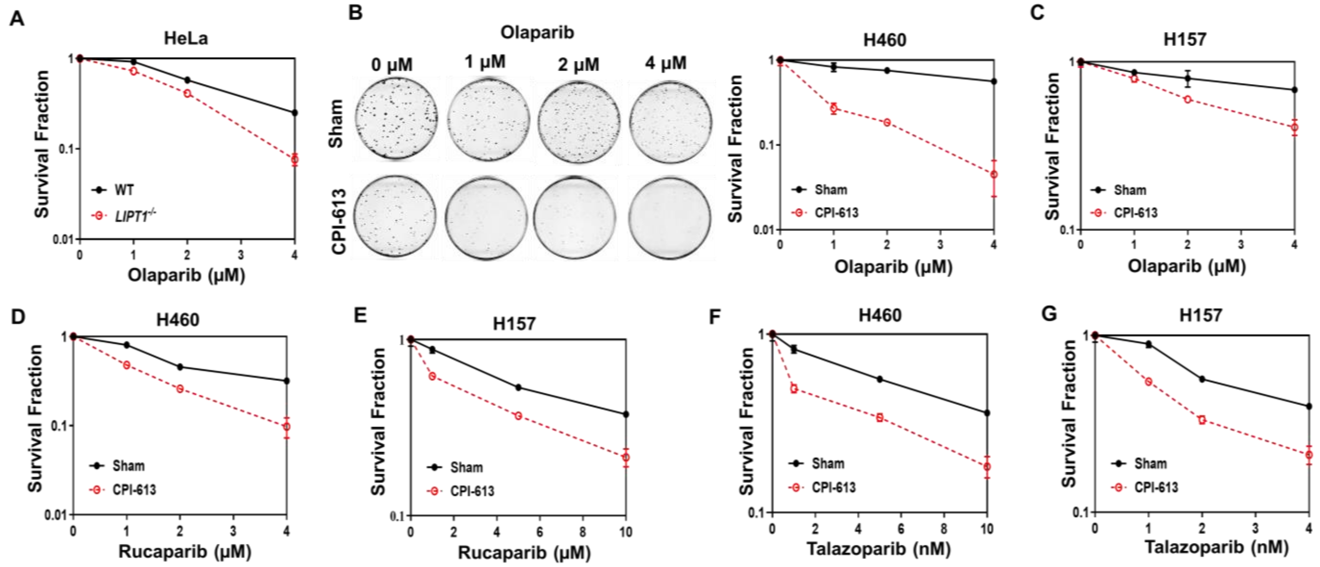


**Fig. S4.** (**A**) Clonogenic survival of WT and *LIPT1*$^{-/-}$ HeLa cells following indicated doses of olaparib. (**B-G**) Clonogenic survival of H460 (B, D, F) and H157 (C, E, G) cells treated with or without 100 μM CPI-613, in combination with indicated doses of olaparib (B, C), rucaparib (D, E) and talazoparib (F, G). Survival fractions were normalized to the DMSO-treated cells.

**Supplemental Figure S5**

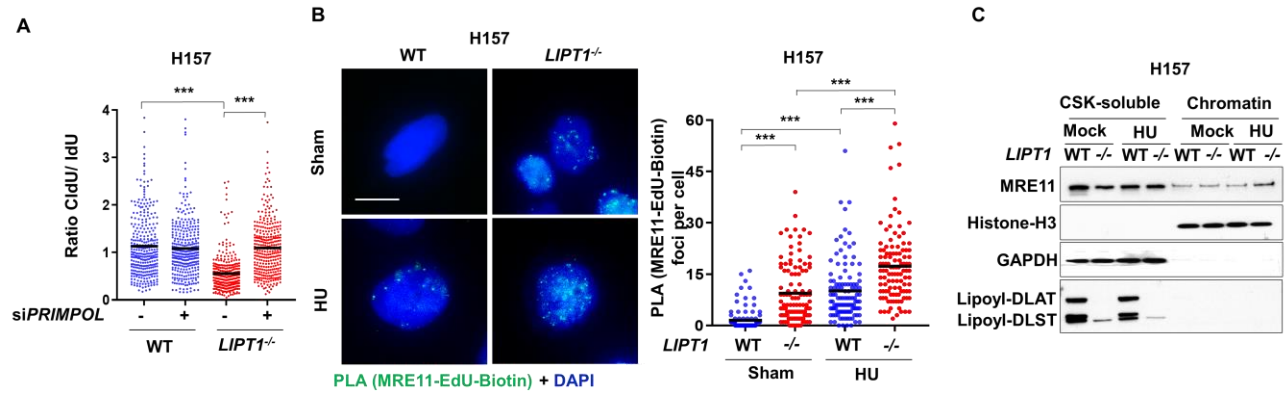


**Fig. S5.** (**A**) CIdU/IdU ratios in WT and *LIPT1*$^{-/-}$ H157 cells after *PRIMPOL* knockdown. Cells were sequentially labeled with IdU (20 min) and CldU (20 min), then treated with HU (4 mM, 4 hours), followed by DNA fiber assay. (**B**) Representative images and quantification of MRE11 by SIRF in WT and *LIPT1*$^{-/-}$ H157 cells. Scale bars, 10 µm. (**C**) Immunoblot of MRE11 in soluble (GAPDH marker) and chromatin-bound (γH2AX, histone H3 markers) fractions from WT and *LIPT1*$^{-/-}$ H157 cells ± HU (2 mM, 2 hours).
